# Multifaceted Changes in Synaptic Composition and Astrocytic Involvement in a Mouse Model of Fragile X Syndrome

**DOI:** 10.1101/615930

**Authors:** Anish K. Simhal, Yi Zuo, Marc M. Perez, Daniel V. Madison, Guillermo Sapiro, Kristina D. Micheva

## Abstract

Fragile X Syndrome (FXS), a common inheritable form of intellectual disability, is known to alter neocortical circuits. However, its impact on the diverse synapse types comprising these circuits, or on the involvement of astrocytes, is not well known. We used immunofluorescent array tomography to quantify different synaptic populations and their association with astrocytes in layers 1 through 4 of the adult somatosensory cortex of a FXS mouse model, the FMR1 knockout mouse. The collected multi-channel data contained approximately 1.6 million synapses which were analyzed using a probabilistic synapse detector. Our study reveals complex, synapse-type and layer specific changes in the neocortical circuitry of FMR1 knockout mice. In layers 1 and 2/3, there is a significant decrease in the density of excitatory glutamatergic synapses, both VGluT1 and VGluT2 type, and their association with astrocytes, while the changes in inhibitory GABAergic synapses are less pronounced. Meanwhile in layer 4, there is a significant increase in the density of small VGluT1 synapses, with no changes in the astrocytic association of synapses. The ability to dissect the circuit deficits by synapse type and astrocytic involvement, will be crucial for understanding how these changes affect circuit function, and ultimately define targets for therapeutic intervention.

## Introduction

Fragile X Syndrome (FXS) is the most common inheritable form of intellectual disability, affecting approximately 1 in 7,000 males and 1 in 11,000 females across all races and ethnic groups [1]. FXS patients display a wide spectrum of phenotypes, including moderate to severe intellectual disability, autistic behavior, macroorchidism, predisposition to epileptic seizures, and facial abnormalities [2, 3, 4]. FXS is caused by the silencing of the FMR1 gene, which encodes the Fragile X Mental Retardation Protein (FMRP). FMRP is known to play an important role in translation, trafficking, and targeting of a large number of mRNAs in neurons [5, 6, 7]. FMRP also binds to many proteins, suggesting its involvement in a wide variety of functions, such as genome stability regulation, cell differentiation, and ion channel gating [8]. Because FMRP participates in a multitude of processes in cells, it has proven difficult to understand how FMRP deficiency affects the synapses and neuronal circuits in brain to cause the FXS pathology.

The mouse model of this disease, the FMR1 knockout mice, display similar phenotypes to human FXS, such as deficiency in learning and memory [9, 10, 11], sensory processing [12, 13], and social behaviors [14, 15]. However, despite these profound neurological and behavioral deficits, the reported changes at synapses have been rather subtle, with basic synaptic neurotransmission seemingly unaffected. At the synaptic functional level, FMR1 KO mice display region-specific deficits in plasticity, such as abnormal long-term potentiation (LTP) and long-term depression (LTD) [8]. Many of the molecular signaling pathways at synapses appear dysregulated, but the changes are often region and neuron-type specific, and the contributions of specific signaling pathways to the Fragile X pathology have been difficult to untangle [16]. At the synaptic structural level, the most obvious difference is the higher density of immature, long and thin dendritic spines of pyramidal neurons in the cortex of adult FMR1 KO mice compared to WT controls [11, 12, 13, 14, 15].

The unusually long and thin spines which are also found in fixed tissues of FXS patients [9, 10, 17, 18], are similar to the immature spines observed during development [19, 20, 21]. This observation has led to a popular hypothesis that the absence of FMRP in the nervous system causes a defect in spine maturation and pruning, which in turn alters synaptic connectivity and ultimately results in behavioral defects [6, 7, 22, 23, 24]. While dendritic spine morphology and structural dynamics are good indicators of modifications in synaptic connectivity [25, 26, 27], they cannot fully represent the diversity of cortical synapses. For example, the majority of inhibitory synapses terminate on dendritic shafts and somata, and are thus not accounted for by changes at spines. Among the excitatory synapses terminating on spines there are cortico-cortical synapses containing the vesicular glutamate transporter VGluT1 and thalamocortical synapses containing VGluT2 [28, 29], which have very different functions in the cortical circuitry. The impact of FXS is likely to dependent on synapse type because of the differential expression of FMRP across neuronal types [30]. Indeed, a recent study using highly multiplexed array tomography showed the varied impact of FXS on synaptic populations of cortical layer 4 and 5 inFMRO1 KO mice [31].

To add a further layer of complexity, FXS may also affect certain non-neuronal cells. As the most abundant glial cells in the mammalian brain, astrocytes modulate synaptic structure and function [32] and are implicated in many neurodevelopmental diseases [33]. In the mouse brain, astrocytes also express FMRP [34], and FMR1 KO mice have fewer hippocampal synapses associated with astrocytes [35]. Interestingly, astrocyte-specific deletion of FMR1 leads to significantly more immature spines in the mouse motor cortex due to overproduction of spines during development [36]. Whether such astrocytic contribution varies according to synapse type is not yet known.

To better understand the synapse type-specific effects of FXS on the neocortical synaptic circuitry, we investigated the changes in different synaptic populations and their association with astrocytes in the adult mouse somatosensory cortex, an area in which a variety of deficits have been reported for FMR1 KO mice [37, 38, 39]. We focused on the superficial cortical layers where live-imaging studies have revealed changes in dendritic spine formation and turnover [37, 40], but the synapse type specificity of the FXS effects is unknown. In order to investigate large numbers of synapses of different types, we used immunofluorescent array tomography (IF-AT) which allows for the light level detection of individual synapses within brain tissue, and the ability to apply multiple markers to distinguish synapse types [41, 42]. Synaptic density was quantified using automatic synapse detection methods previously developed by our group [43, 44]. Our results reveal multifaceted changes in the composition and astrocytic involvement in the synaptic circuitry of the somatosensory cortex of adult FMR1 KO mice.

## Methods

### Overview

The methods section is divided into two main components — data generation and computational analysis. The data generation section specifies the types of mice used, the antibodies used, and the imaging methodology. The computational analysis section highlights the methods used to automatically analyze the array tomography data, including the detection of synapses by their specific type and the detection of astrocytes.

### Data generation

The datasets investigated were obtained from the somatosensory cortex of adult mice and represent layers one through four. The somatosensory cortex was chosen because of the well-documented deficits in FMR1 KO mice in this cortical region[37, 38, 39]. We focused on the superficial cortical layers for which more information is available through live imaging studies [37, 40]. The average dataset volume was 33, 897*µm*^3^ with a standard deviation of 9, 606*µm*^3^.

### Animals

FMR1 KO mice were obtained from Dr. Stephen T. Warren, Emory University. Thy1-YFP-H mice were purchased from JAX. All mice were backcrossed with C57BL/6 mice more than 10 generations to produce congenic strains. For the current experiments, YFP+ WT males were crossed with YFP-FMR1+/- females, and only male offspring littermates were used for the experiments. WT mice refer to FMR1+/y, and KO mice are FMR1-/y. Because the YFP expression was highly variable between animals, we did not use it in the analysis. The mice were four months old when they were sacrificed. Further details about the mice are in Supplemental Table S1.

### Array tomography

The tissue was prepared using standard array tomography protocols [42]. Mice were group-housed in the UCSC animal facility, with 12 hour light-dark cycles and access to food and water ad libitum. All procedures were performed in accordance with protocols approved by the Animal Care and Use Committee (IACUC) of UCSC. The mice were anesthetized by halothane inhalation and their brains quickly removed, cut into 2*mm* slices, fixed by immersion in 4% paraformaldehyde in phosphate-buffered saline (PBS) for 1 hour at room temperature, then left in the fixative overnight at 4 °C. After rinsing in PBS, the somatosensory cortex was dissected out, quenched in 50*mM* glycine in PBS for 30 minutes and dehydrated in a series of ethanol washes (50%, 70%, 70%) at 4 °C, then infiltrated and embedded in LRWhite resin in gelatin capsules, and polymerized at 50 °C for 24 hours.

To prepare ribbons of serial sections, the blocks were trimmed around the tissue to the shape of a trapezoid, and glue (Weldwood Contact Cement diluted with xylene) was applied with a thin paint brush to the leading and trailing edges of the block pyramid. The embedded plastic block was cut on an ultramicrotome (Leica Ultracut EM UC6) into 70*nm*-thick serial sections, which were mounted on gelatin-coated coverslips.

### Immunolabeling

Sections were processed for standard indirect immunofluorescence, as described in [42]. Antibodies were obtained from commercial sources and are listed in Table 1. Array tomography specific controls are presented in Supplemental Table S2. The sections were incubated in 50 mM glycine in TBS for 5 minutes, followed by blocking solution (0.05% Tween-20 and 0.1% BSA in TBS) for 5 minutes. The primary antibodies were diluted in blocking solution as specified in Table 1, and were applied for 2 hours at room temperature or overnight at 4 °C. After a 15 minutes wash in TBS, the sections were incubated with Alexa dye-conjugated secondary antibodies, highly cross-adsorbed (Life Technologies), diluted 1:150 in blocking solution for 30 minutes at room temperature. Finally, sections were washed with TBS for 15 minutes, rinsed with distilled water and mounted on glass slides using SlowFade Gold Antifade Mountant with DAPI (Invitrogen). After the sections were imaged, the antibodies were eluted using a solution of 0.2 M NaOH and 0.02% SDS for 20 minutes, and new antibodies were reapplied. Several rounds of elution and re-staining were applied to create a high-dimensional immunofluorescent image. Samples were immunostained side by side in pairs, consisting of one WT and one KO sample, and imaged immediately after completion of staining.

**Table 1.** Antibodies used for the experiments.

| Antigen | Host | Antibody Source | RRID | Dilution |
| --- | --- | --- | --- | --- |
| Synapsin | Rabbit | Cell Signaling 5297 | RRID:AB_2616578 | 1:100 |
| PSD95 | Rabbit | Cell Signaling 3450 | RRID:AB_2292883 | 1:100 |
| VGluT1 | Guinea pig | Millipore AB5905 | RRID:AB_2301751 | 1:5000 |
| VGluT2 | Guinea pig | Millipore AB2251 | RRID:AB_2665454 | 1:5000 |
| GAD2 | Rabbit | Cell Signaling 5843 | RRID:AB_10835855 | 1:100 |
| Gephyrin | Mouse | NeuroMab 75-443 | RRID:AB_2636851 | 1:100 |
| Glutamine synthetase | Mouse | BD Biosciences 610517 | RRID:AB_397879 | 1:25 |

### Imaging method

The immunostained ribbons of sections were imaged on an automated epifluorescence microscope (Zeiss AxioImager Z1) using a 63× Plan-Apochromat 1.4 NA oil objective. To define the position list for the automated imaging, a custom Python-based graphical user interface, MosaicPlanner (obtained from https://code.google.com/archive/p/smithlabsoftware/), was used to automatically find corresponding locations across the serial sections. Images from different imaging sessions were registered using a DAPI stain present in the mounting medium. The images from the serial sections were also aligned using the DAPI signal. Both image registration and alignment were performed with the MultiStackReg plugin in FIJI [45].

### Computational analysis

A main goal of this analysis is to examine the effects of the lack of FMRP protein on the synaptic composition of the somatosensory cortex. This requires the ability to quantify synapses by their molecular composition and their adjacency to an astrocytic process. To achieve this, we took existing methods and expanded their scope to meet the computational challenges posed by these experiments, including developing a method for detecting astrocyte processes adjacent to synapses.

### Synapse detection

For the present purposes, we define ‘synapse type’ as a specific combination of synaptic proteins. For example, a GABAergic (inhibitory) synapse type is defined by the presence of the general presynaptic marker, synapsin; the postsynaptic marker of inhibitory synapses, gephyrin; and the presynaptic marker of inhibitory synapses, GAD. A glutamatergic (excitatory) synapse is defined by the presence of the general presynaptic marker, synapsin, and the postsynaptic marker for excitatory synapses, PSD-95. A glutamatergic synapse with VGluT2 and adjacent to an astrocytic process is defined by the presence of synapsin, VGluT2, PSD-95, and GS, a marker for astrocytes.

Detecting synapses by their molecular composition is the first step of the computational pipeline. In order to quantitatively analyze large array tomography volumes, it is vital to find an appropriate synapse detection technique. The majority of published synapse detection methods use traditional machine learning approaches [46, 47, 48, 49]. These approaches all consist of a few common steps to detect synapses. First, for each synapse type, a large number of synapses are manually identified and labeled in the array tomography data. Next, a classifier (such as a support vector machine or convolutional neural network) is trained with these manual annotations. Lastly, the entire dataset is appropriately parcellated and potential synapses are labeled by the classifier. While this method works well for certain questions in synapse biology, the difficulty in manually labeling different synapse types in immunofluorescent data renders it ineffective for our applications.

The probabilistic synapse detection method introduced in [43], is a synapse type focused approach which does not require any training data, making it a viable option for exploring synapses imaged via array tomography. ‘Synapse type’ focused means the user specifies the molecular composition and the relative spatial arrangement along with the size of the synaptic markers prior to running the probabilistic synapse detection method. The combination of a user-defined synapse type and marker size is called a ‘query,’ as highlighted in Figure 1.

**Figure 1.**
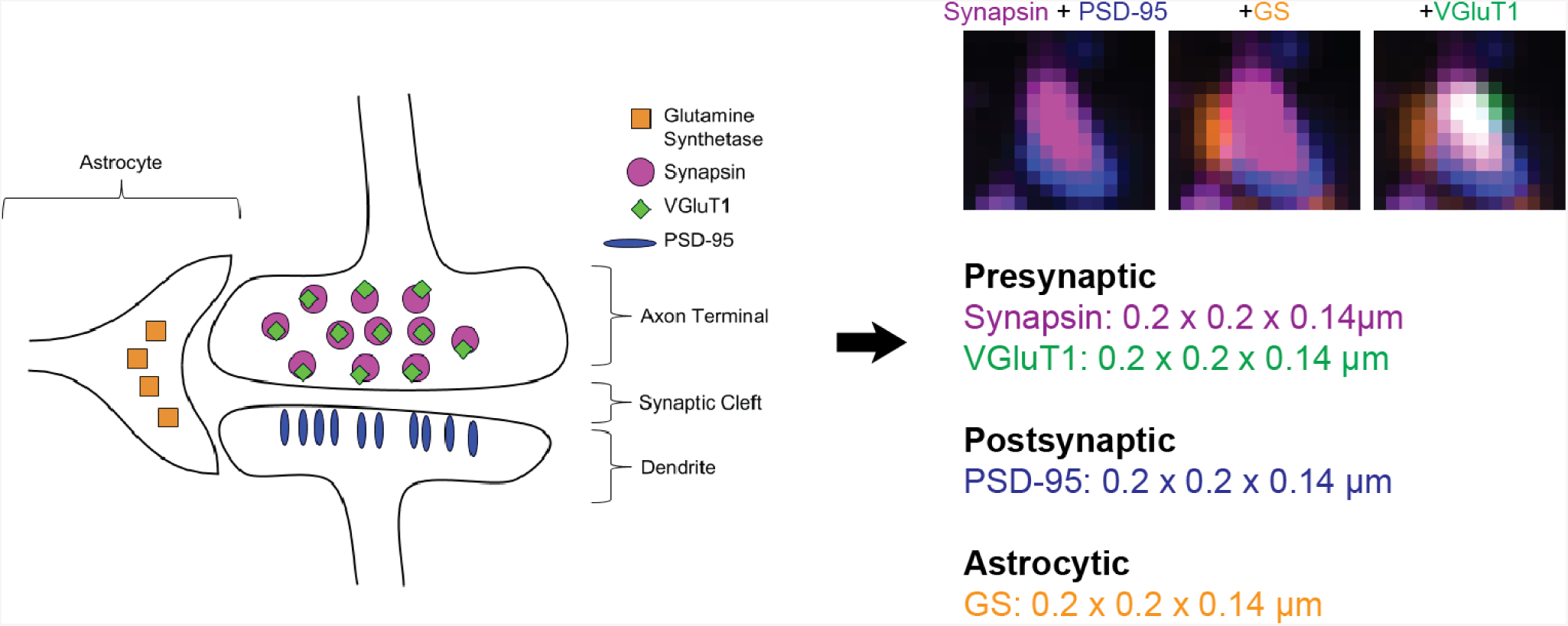
Outline of a query. The cartoon on the left side shows the relative spatial arrangement of the different fluorescent markers used to detect an excitatory synapse expressing VGluT1, next to an astrocyte process. This visual description of a synapse is translated into a query, shown to the right of the large blue arrow. A query is a user-defined description of what the synapse type of interest should ‘look’ like. In this case, the presynaptic protein markers - synapsin and VGluT1, are expected to colocalize (occupy the same 3D space) with each other. Furthermore, the presynaptic, postsynaptic, and astrocyte markers (as a group) are all expected to be next to each other. The top right portion of the figure shows three 1.5 × 1.5*µm* cutouts of different marker combinations showing what the query looks like in the data. The first cutout shows the synapsin and PSD-95 punctum overlaid; the second cutout includes the GS punctum and the third cutout includes the VGluT1 punctum.

A query to detect a glutamatergic synapse would look like the following: a PSD-95 punctum of a minimum of 2*px* × 2*px* × 2*slices* (which is 0.2*µm* × 0.2*µm* × 0.14*µm* for our data) adjacent to a synapsin punctum of the same size. Adjacency in this case means that the puncta of the two different antibody markers do not occupy the same space but instead are juxtaposed from each other. Since PSD-95 is a postsynaptic protein and synapsin is presynaptic protein, this simple glutamatergic synapse query follows the known biological model for a glutamatergic synapses. In [43], the query comprises only of presynaptic and postsynaptic makers. In this work, we have expanded the query to comprise of presynaptic, postsynaptic, and astrocytic markers. This combination of markers is often referred to as a ‘tripartite synapse’ [50]. As the left side of Figure 1 shows, the tripartite synapse model assumes that the molecular markers for each ‘subclass’ (presynaptic, postsynaptic, astrocytic) lie adjacent relative to each other.

In summary, the probabilistic synapse detector is a method of detecting specific synapse types. Instead of requiring the user to manually annotate multiple instances of a synapse type to train a machine learning classifier, the query-based approach asks the user to define a synapse by specifying basic characteristics, that is, the requisite markers, requisite punctum volume for each marker (which depends on the microscope known resolution), and their relative spatial arrangement.

Once the query has been established, the probabilistic synapse detection method follows the query to automatically detect synapses matching the query. Figure 2 shows an example pipeline going from the raw input data to the result probability map. The details of the synapse detection method are described in [43] and its applications to antibody characterization are studied in [44]. Both report extensive validation, indicating that the tool is ready to address the novel biological questions in this work.

**Figure 2.**
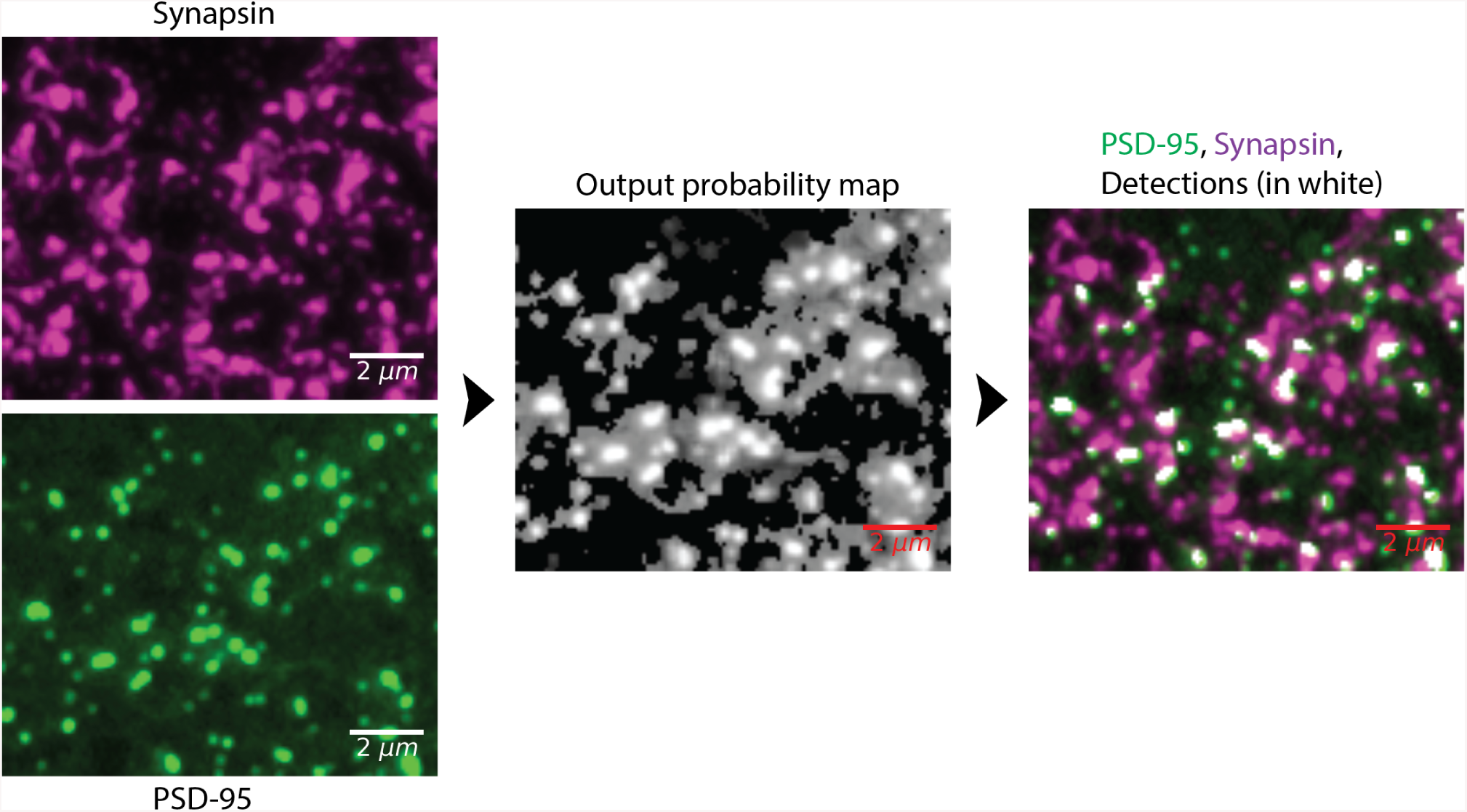
Probabilistic synapse detection pipeline. The first column shows the raw PSD-95 and synapsin data. The second column shows the output of the synapse detection method, where the value at each pixel is the probability that pixel belongs to the specified definition of a synapse. The third column shows the detections (in white) overlaid upon the superposition of the PSD-95 and synapse data. For this visualization, the definition of a synapse was the adjacency of a PSD-95 and synapsin punctum of a minimum size of 0.2*µm* × 0.2*µm* × 0.07*µm* which corresponds to 2*px* × 2*px* × 1*slice*.

### Synapse type definitions

For the analysis presented in this work, we used the queries listed in Table 2. A synapse of a particular type is defined as having the relevant markers, with all the markers being of the specified size. For this study, we required the markers to span one or more slices, depending on the desired synapse size, and have a minimum *x, y* size of 0.2*µm* × 0.2*µm*. The one exception is the definition of VGluT2 synapses where the VGluT2 marker is required to span two or more adjacent slices. This is due to the properties of the VGluT2 antibody, which in addition to the expected robust label of a synapse subpopulation, also gives higher, randomly distributed, background signal (see Supplemental Table S2 for more information).

**Table 2.**
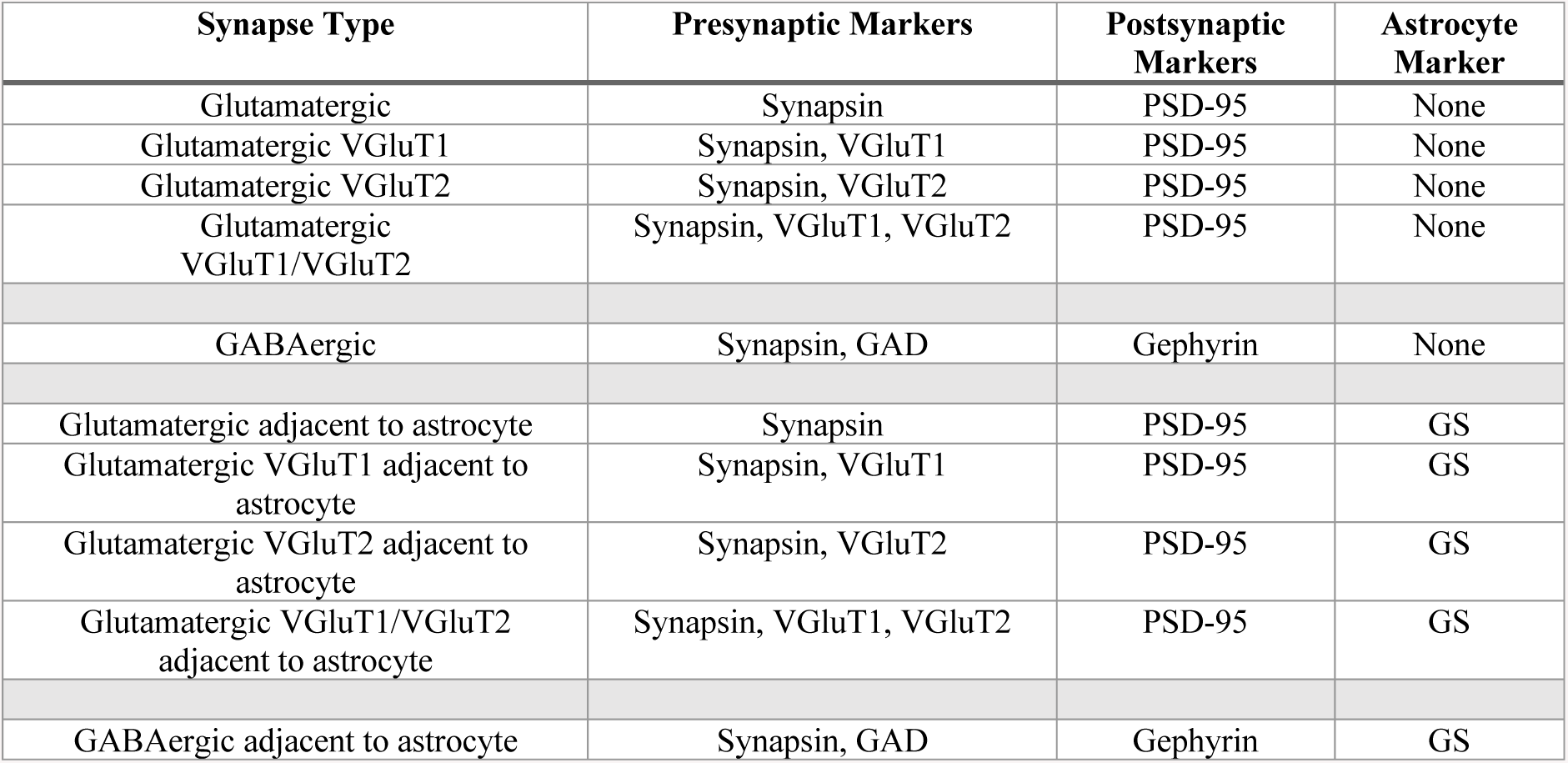
Queries used for this analysis.

Each synapse type was further subdivided in three sizes: small, medium, and large. A small synapse is defined as spanning one slice, a medium synapse is defined as spanning two slices, and a large synapse is defined as spanning three or more slices. To calculate the synapse density of synapses that only span a single slice, a query where synapses span two or more slices is subtracted from a query where synapses span one or more slices. In the same vein, to calculate the synapse density of synapses that span only two slices, a query where synapses span three or more slices is subtracted from a query where the synapses span two or more slices. ‘All synapses’ of a type are defined as having punctum markers than span one or more slices.

### Volume calculation

Synapse density is calculated as the number of synapses detected in a dataset over the volume of the dataset. The volume of the dataset is defined by the volume of the neuropil (i.e., excluding the volume occupied by cell nuclei and large blood vessels, which can vary significantly between areas). When choosing the areas to image, we avoided large blood vessels. To calculate the volume of cell nuclei, the nuclear stain DAPI was converted into probability space using the methods outlined in [44]. Briefly, the value at each pixel in probability space is the probability it belongs to the foreground, with a range of 0 to 1. To do so, the background noise is modeled and the foreground probability is one minus the probability the pixel belongs to the background. The background noise is modeled as a Gaussian, for which the mean and variance is calculated from the raw data itself. Once the probability map is calculated, it is thresholded (t=0.6, chosen by observation) and cleaned up by a sequence of morphological operations. The code is available for download on the project’s website. In summary, the volume of the neuropil was obtained by subtracting the volume of the DAPI stained nuclei from the total imaged volume.

### Colocalization analysis

To examine the spatial relationships between glial and postsynaptic markers, we used the Colocalization test in Fiji [45], applying the van Steensel method of randomization [51]. For each pair of channels, we used a 40 × 26*µm* region of interest through the image stack. The excitatory and inhibitory presynaptic/postsynaptic marker pairs, VGluT1/PSD95 and GAD/Gephyrin were used for comparison.

### Statistical analysis

Statistical analysis to determine significance between the two populations was done via a two-tailed unpaired t-test, shown to be applicable to very small sample sizes [52], such as used in this study (n=4). Because of the potential problems with any small sample statistical test, we are also presenting the data from each individual sample in the supplemental figures section. Plots showing the data for each synapse type, size, and layer are found in Supplemental Figures S1, S2, S3, S4, S5, S6.

### Data and code availability

The code and raw data are available for download at https://aksimhal.github.io/astrocytes-synapses-fxs/.

## Results

We used immunofluorescent array tomography (IF-AT) to quantify the synaptic density, composition and glial involvement in layers 1 through 4 of the somatosensory cortex of adult FMR1 knockout (KO) mice and wild-type (WT) mice. IF-AT is based on digital reconstruction of images acquired from arrays of serial ultrathin sections (70 nm) attached to coverslips, immunofluorescently labeled and imaged under a fluorescence microscope. The use of ultrathin sections allows the light level detection of individual synapses, while the possibility of applying multiple immunofluorescent markers (10 or more) enables the identification of different synaptic populations [41]. Synaptic density was quantified using automatic synapse detection methods previously developed by our group [43, 44]. Besides the already published validation of our method, results from WT mice were also compared to available estimates in the literature as an additional control.

### Overview of the datasets and detected synapses

Volumes of approximately 140 × 400 × 3*µm* spanning layers 1 through 4 of the somatosensory cortex of FMR1 KO mice and WT mice were imaged, as shown in Figure 3. We detected an average of 200,000 synapses in each volume for a total of approximately 1.6 million synapses across all eight datasets. Excitatory synapses were identified by the presence of immunofluorescent signals from both synapsin, a presynaptic protein, and PSD-95, a protein of the postsynaptic scaffold of excitatory synapses. Excitatory synapses were further subdivided depending on their vesicular glutamate transporters into VGluT1 positive, generally thought to be of intracortical origin, and VGluT2 positive, belonging predominantly to thalamocortical inputs [28, 29, 53]. Inhibitory synapses were identified by the presence of the general presynaptic marker synapsin and the presynaptic marker for GABAergic synapses, glutamic acid decarboxylase (GAD), together with the postsynaptic marker gephyrin. Astrocytes, including their processes, were detected using an antibody against glutamine synthetase (GS) [54, 55], which allowed for the identification of the fraction of synapses that are immediately adjacent to astrocytic processes. In addition to identifying synapses based on combinations of different markers, synapses were also analyzed based on their size, because size is known to correlate with the maturity and strength of a synaptic connection. Newly formed synapses tend to be small, and at mature synapses the size of the postsynaptic density is known to correlate well with synaptic strength [56]. Small synapses were identified as having the relevant markers on only one slice, medium synapses — on two consecutive slices, and large synapses — on three or more consecutive slices. Visual inspection of the datasets did not uncover any obvious differences in immunofluorescence intensity and pattern for any of the markers between the KO and WT mice. The cortical thickness was also comparable between the two conditions (0.88 ± 0.03*mm* for KO vs. 0.89 ± 0.02*mm* for WT, *p* = 0.71).

**Figure 3.**
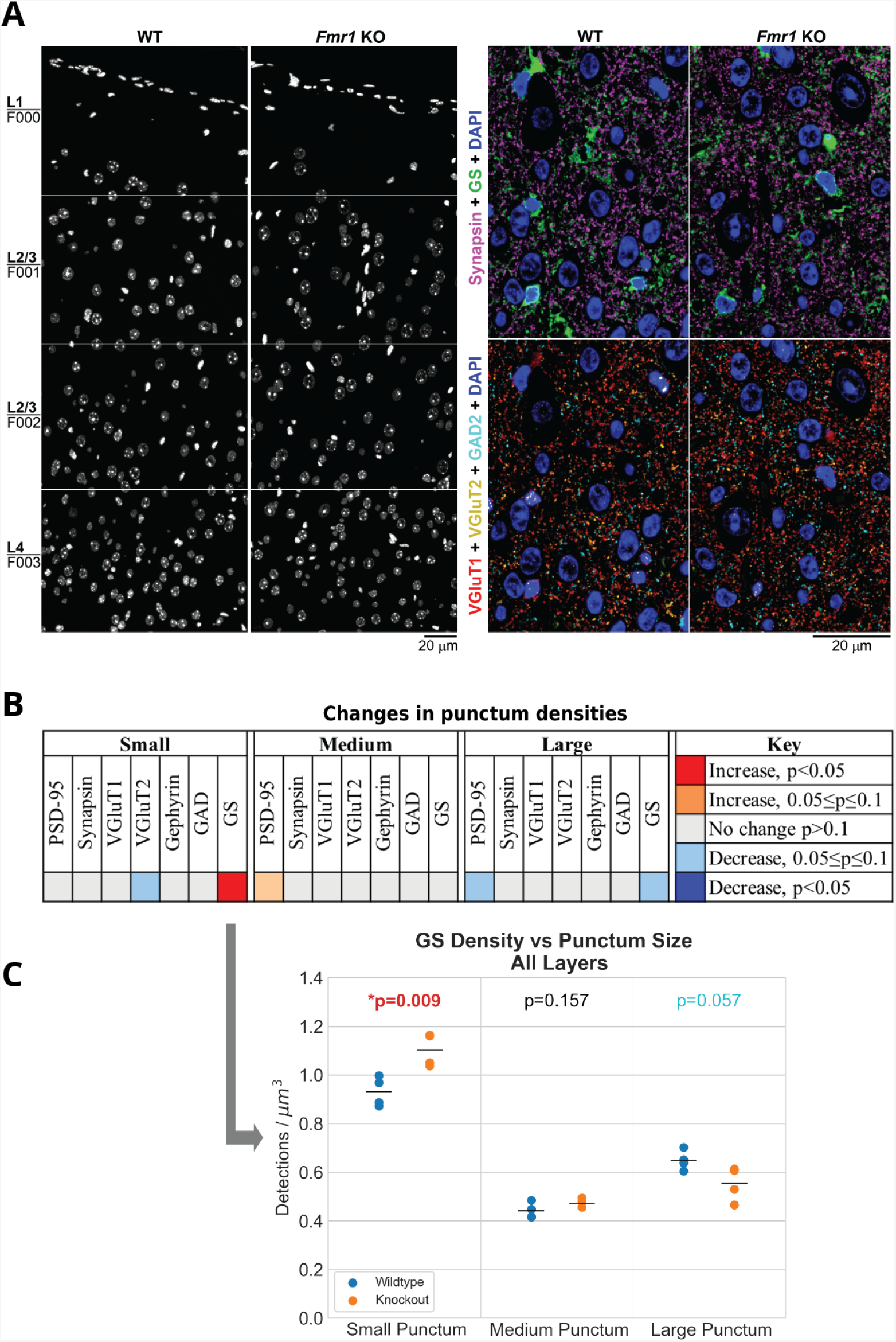
Overview of the datasets. A, Immunofluorescent array tomography of wild-type and FMR1 knockout mouse somatosensory cortex. The left panel shows the imaged area in each sample, consisting of four tiles spanning cortical layers 1 through 4. DAPI staining of nuclei, volume reconstruction of 30 serial sections, 70 nm each. The right panel shows the immunofluorescence for synaptic (synapsin, VGluT1, VGluT2 and GAD2) and glial (GS) markers in wild-type and knockout mouse somatosensory cortex layer 4, volume reconstruction of 10 serial sections, 70 nm each. B, Summary of single channel punctum density changes between wild-type and knockout mice with all layers averaged. C, This plot shows the density distribution of GS puncta by size. ‘Small’ puncta span one slice, ‘medium’ puncta span two slices, and ‘large’ puncta span three or more slices.

### Single channel analysis

Quantification of the puncta density of the different synaptic markers did not reveal any statistically significant differences (*p* > 0.05) when averaging Layers 1-4, as shown in Figure 3B and in more detail in Supplemental Figure S1. There was a tendency for a decrease in the number of small VGluT2 puncta (*p* = 0.065) and large PSD95 puncta (*p* = 0.065). On the other hand, medium PSD-95 puncta tended to increase (*p* = 0.056), as shown in the middle panel of Figure 3.

The only significant change that was detected was in the density of GS puncta of different sizes: in FMR1 KO mice there were more small puncta (*p* = 0.01), as well as a tendency for a decrease in the number of large GS puncta (*p* = 0.07), as shown in the bottom panel of Figure 3.

Even though we did not detect any significant changes in the densities of puncta of the different synaptic markers, this does not necessarily preclude changes in the synaptic populations of FMR1 KO mice. The synaptic proteins assessed in our study are indeed highly enriched at synapses, but they are also found at extra-synaptic sites, and thus their presence does not necessarily equate to the presence of a synapse.

### Overall synapse densities

A much more accurate detection of synapses is achieved by using combinations of synaptic makers, ideally at least one presynaptic and one postsynaptic marker, as specified by our synapse detection algorithm [43, 44]. Indeed, using such combinations of synaptic markers, the detected synapse densities and distributions in WT mice are consistent with previous estimates as shown in Figure 4. The overwhelming majority of cortical synapses are known to be either excitatory glutamatergic or inhibitory GABAergic synapses [57]. Thus, the total density of synapses was estimated by the sum of the densities of the detected glutamatergic (synapsin + PSD95 markers) and GABAergic synapses (synapsin + GAD + gephyrin markers) resulting in approximately 1.94 synapses per *µm*^3^ of embedded tissue. Because tissue dehydration and embedding with our protocol results in approximately 23% linear shrinkage, or 54% volumetric shrinkage [46], this equals to 0.9 synapses per *µm*^3^ of unprocessed tissue, very similar to the reported synapse density in mouse cortex [58].

**Figure 4.**
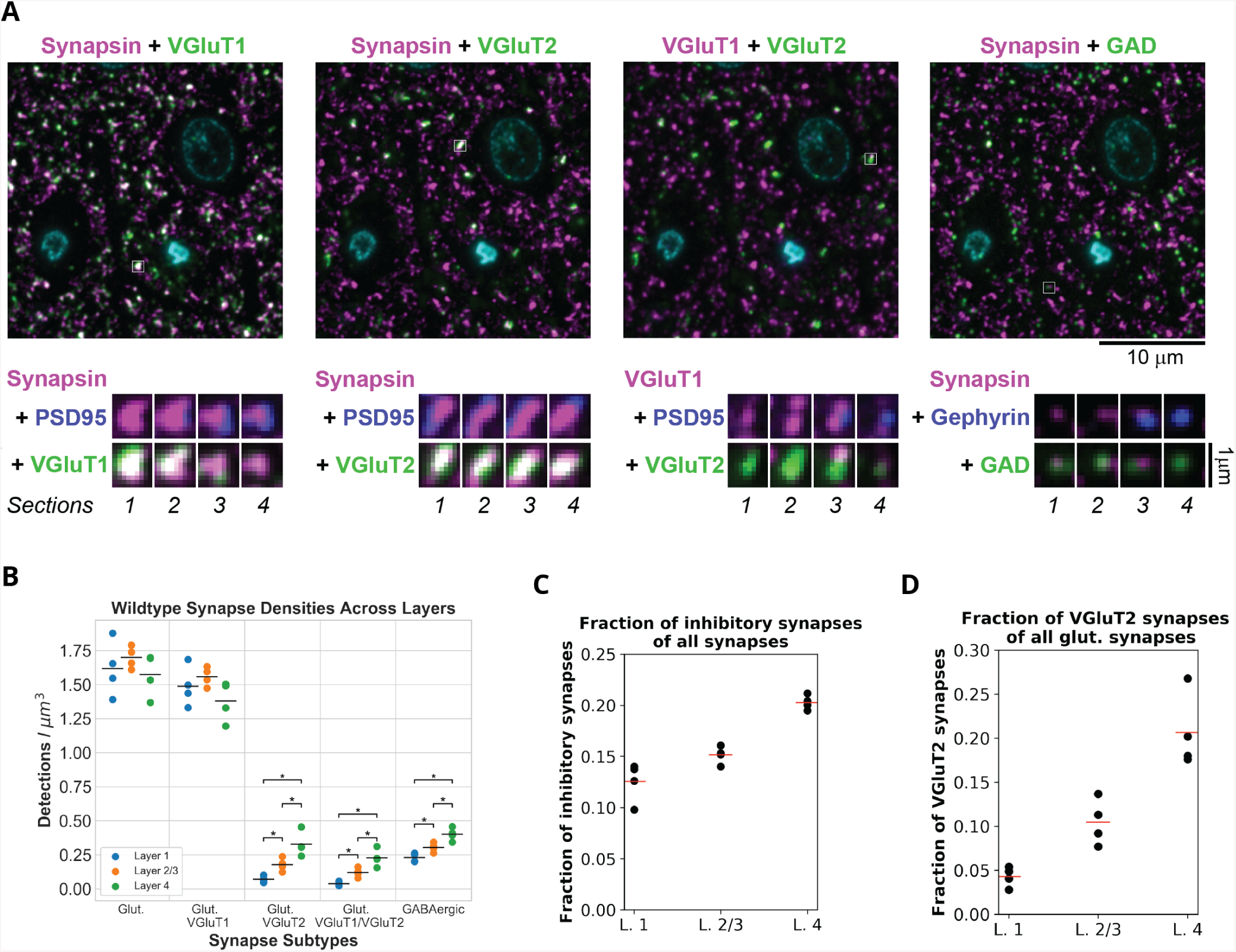
Overview of wild-type synapse density distributions. A, top. Multiplexed immunofluorescence from the same area of a single section of the somatosensory cortex of a WT mouse. The nuclear stain DAPI is in cyan. White squares highlight individual synapses, which in A, bottom, are shown at higher magnification on 4 consecutive serial sections. B, The distribution of synapse types across different layers. Layers with significant differences for a synapse type are marked. While there is no significant difference in layer densities for glutamatergic synapses overall and for glutamatergic synapses with VGluT1, there is a significant difference between layers for glutamatergic synapses with VGluT2, with both VGluT1 and VGluT2, and for inhibitory synapses. C, Fraction of inhibitory synapses in layers 1 through 4 of mouse somatosensory cortex. D, Fraction of VGluT2 synapses in layers 1 through 4 of the mouse somatosensory cortex.

The relative contributions of inhibitory and excitatory synapse types that we detect is consistent with the known synapse composition of mouse cortex. The percent of inhibitory GABAergic synapses in our detections varies between 12% in layer 1 to 20% in layer 4 (Figure 4F). Electron microscopy (EM) counts in mouse somatosensory cortex show that inhibitory synapses consist of 11% of the synapses in layer 1 [59] and 18% of the synapses in layer 4 [60].

The layer distribution of VGluT2 synapses is consistent with their known preference for layer 4 (Figure 4G). VGluT2 is known to label thalamocortical synapses which target mostly layer 4 and lower layer 2/3 [61, 62]. Thalamocortical synapses, identified either by degeneration techniques, anterograde transport of lectin, or VGluT2 immunostaining, have been shown to comprise approximately 20% of glutamatergic synapses in layer 4 of mouse somatosensory cortex [63, 64, 65] and we indeed see 21% in layer 4.

To further verify the accuracy of our detections, we used a different way to calculate the density of excitatory glutamatergic synapses. These synapses can be subdivided into two major populations depending on the vesicular glutamate transporters (VGluTs) at the presynaptic site, with the majority of excitatory synapses containing VGluT1, and a smaller population, mostly concentrated in layer 4, VGluT2. In addition, some synapses express both VGluT1 and VGluT2 [66]. Thus, the density of glutamatergic synapses should be approximately equal to the densities of VGluT1 and VGluT2 synapses minus VGluT1/VGluT2 synapses to prevent double counting of the same synapse. This was indeed the case, as seen in Table 3.

**Table 3.** Density distribution of excitatory synapses across layers as calculated by two methods. Units are number of synapses per cubic micron.

|  | Glut | VGluT1+VGluT2-VGluT1/2 |
| --- | --- | --- |
| ***L1*** | 1.617 | 1.629 |
| ***L2/3*** | 1.700 | 1.750 |
| ***L4*** | 1.574 | 1.585 |

### Changes in synaptic densities in FMR1 KO mice

Next, we compared the densities of the different synapse populations in the WT mice to the FMR1 KO mice. Even though the individual synaptic marker puncta did not show any statistically significant differences in the two conditions, there were wide-ranging changes in synaptic densities. These changes were dependent on the synapse type, size, as well as cortical layer, as shown in Figure 5 and in more detail in Supplemental Figure S2. There was an increase of small glutamatergic VGluT1 synapses in layer 4 accompanied by a decrease in large VGluT1 synapses in layers 1 and 4. VGluT2 synapses, on the other hand, showed a rather consistent decrease in density, regardless of their size, with the majority of changes occurring in layers 1 and 2/3, but not layer 4 (except for large VGluT1/2 synapses). Large inhibitory synapses decreased across all layers examined without detected changes in small and medium size inhibitory synapses.

**Figure 5.**
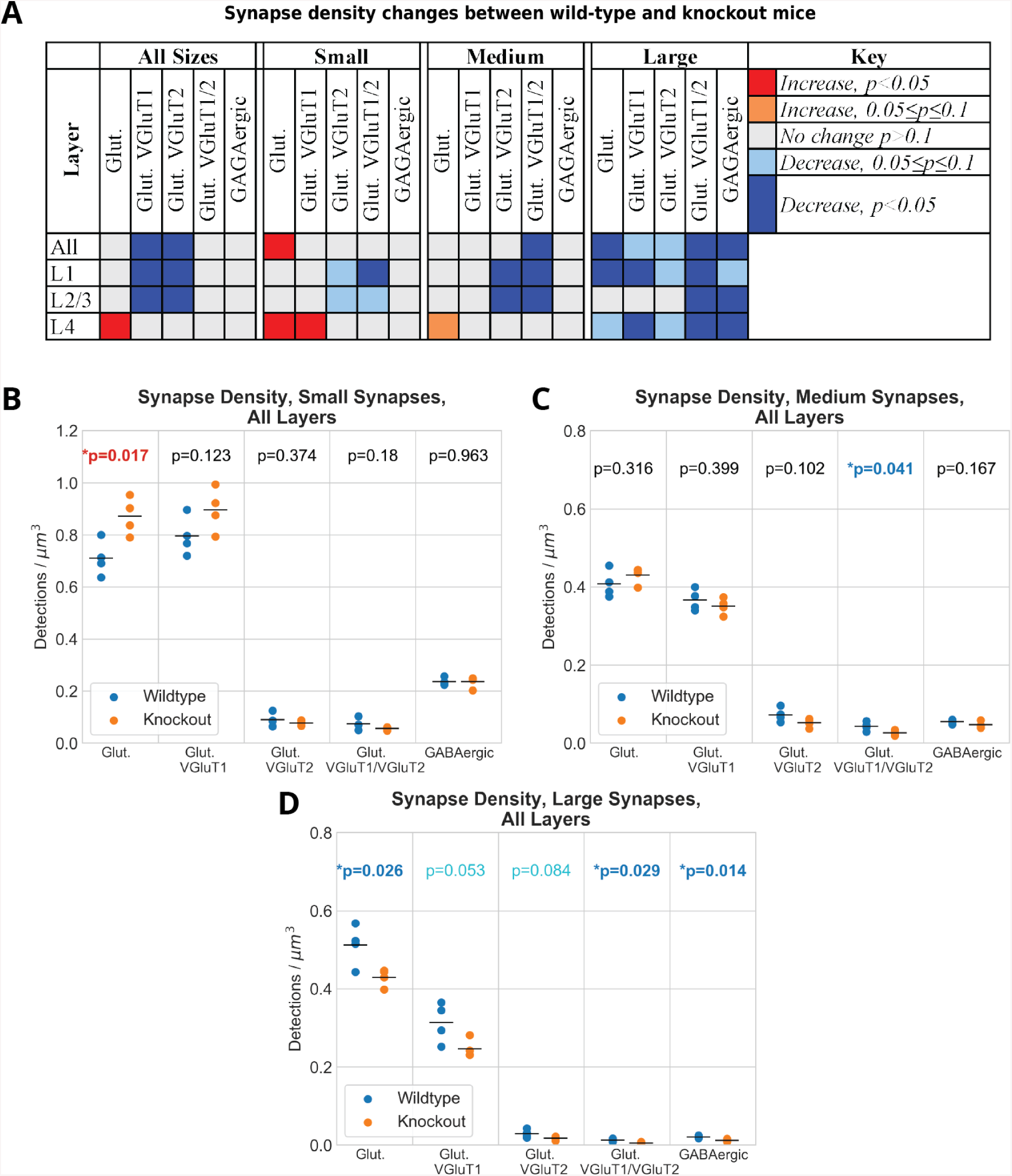
Changes in synapse densities across layers. A, Summary table showing the synapse types that have a significant increase or decrease in synapse density between wild-type and knockout mice. B-D, Differences in synapse density for different synapse types for three expected synapse sizes.

These changes in density of the various synaptic populations resulted in an increase in the excitation-inhibition ratio in the FMR1 KO mice: 6.20±0.56 for the KO vs. 5.24±0.32 in WT (average for layers 1 through 4; *p* = 0.04). Excitation was defined by the number of synapses which contain synapsin and PSD95, and inhibition by the number of synapses containing synapsin, GAD and gephyrin. These changes are highlighted in Figure 6G.

**Figure 6.**
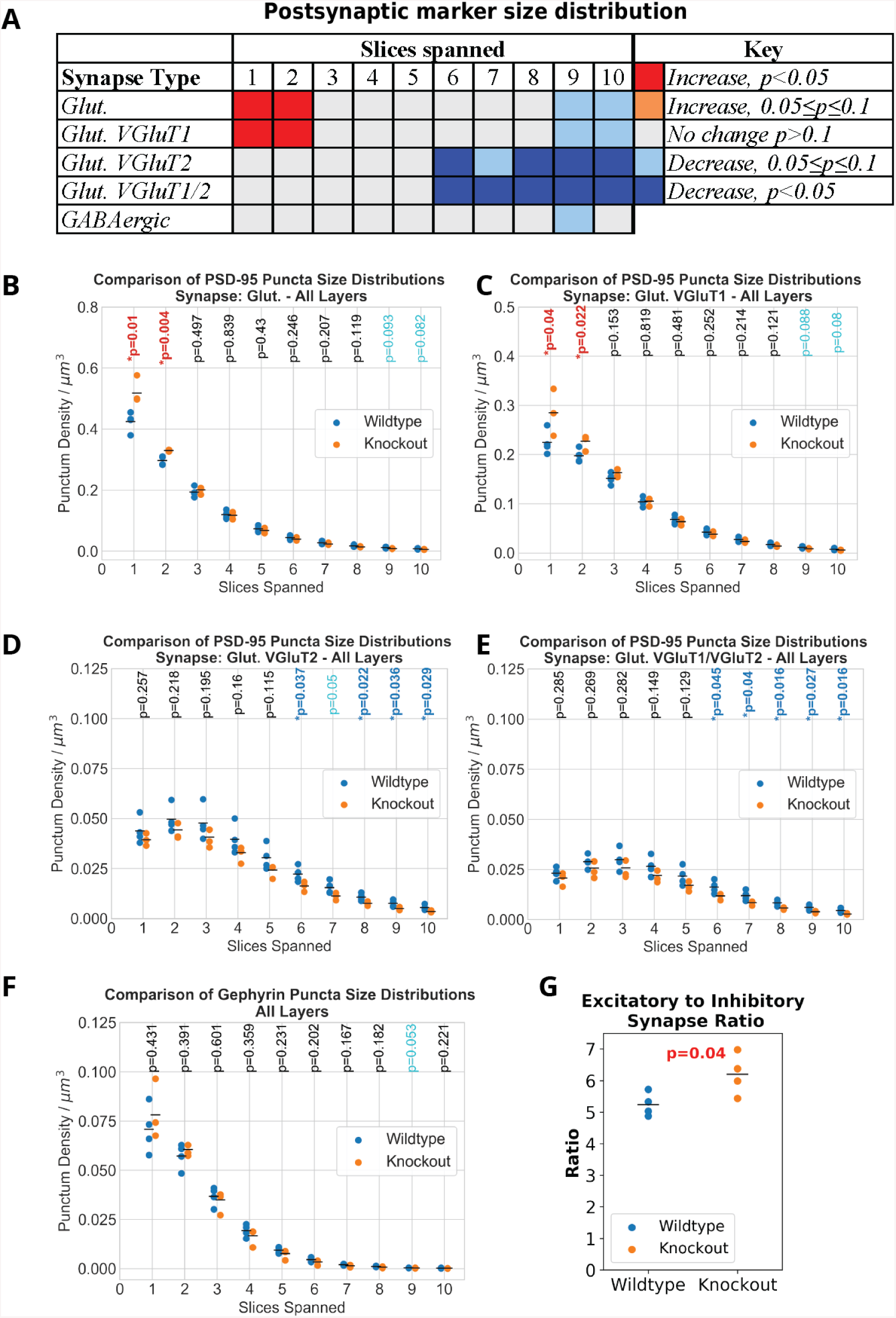
Postsynaptic marker size distributions. A, Summary table showing the differences in the size distributions of PSD-95 puncta associated with synapses by type. The main takeaway is that for excitatory synapses overall, there is an increase in the number of small PSD-95 puncta in FMR1 knockout mice while there is a decrease in the number of very large PSD-95 puncta associated with synapses containing VGluT2. B-E, Plots showing the distribution of PSD-95 puncta for both wild-type and knockout mice. F, Size distribution of gephyrin puncta associated with inhibitory synapses. G, Plot shows the significant increase in the ratio of excitatory to inhibitory synapses in knockout mice.

Because the strength of excitatory synapses is known to be proportional to the size of the postsynaptic density, we also analyzed the changes specifically at the postsynaptic side as shown in Figure 6A-F and in more detail in Supplemental Figure S3. There was a statistically significant increase in the density of glutamatergic and specifically VGluT1 synapses with small PSD-95 puncta (spanning only 1 or 2 slices), and a significant decrease in the density of the VGluT2 synapses with large PSD-95 puncta (spanning 6 or more slices). No changes in the densities of inhibitory synapses depending on the size of gephyrin puncta were detected, as shown in more detail in Supplemental Figure S4.

### Involvement of glia (astrocytes)

Astrocytes are intimately involved in synaptic function and their processes are found adjacent to many synaptic clefts in the neocortex. Because astrocytes in the mouse also express FMRP [34] and are suspected to have a role in FXS pathogenesis [36], we analyzed the potential changes in astrocytic involvement at synapses in the FMR1 KO mice. Astrocytes were detected using antibodies to glutamine synthetase, an enzyme known to be expressed predominantly by this cell type [54, 55] and specifically found at the peripheral astrocytic processes that contact synapses [67, 68]. Figure 7A show an example of the data.

**Figure 7.**
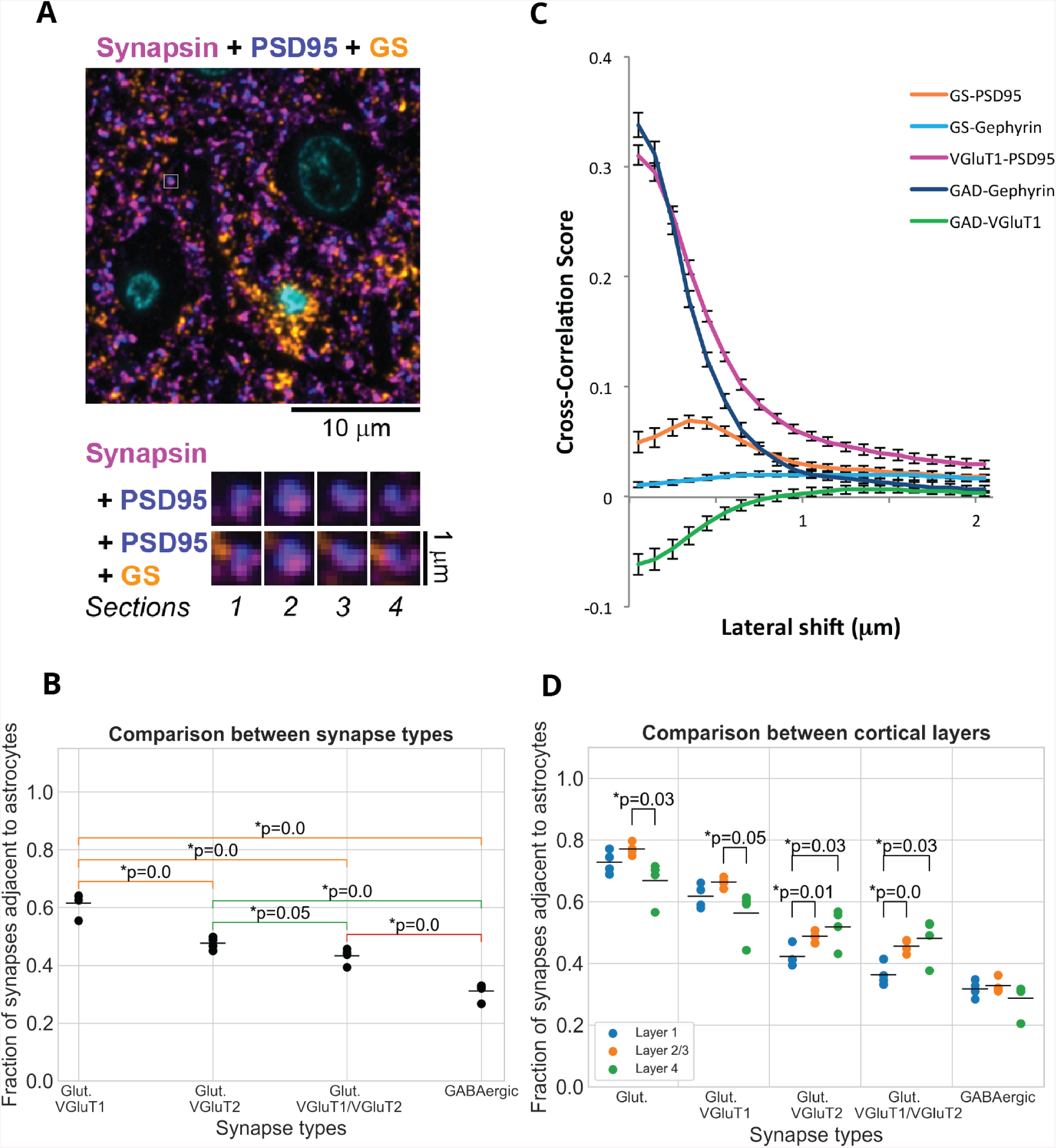
Astrocytic association of synapses in wild-type mice. A, Portion of the wild-type synapse data with a synapse highlighted with a white box. ?Below are serial sections though the highlighted synapse, shown at higher magnification. B, The density of synapses associated with an astrocytic process decreases dramatically for all synapse types across all layers. C, Cross-correlation score as a function of lateral offset between the two channels. The synaptic marker pairs VGluT1 - PSD95 and GAD - Gephyrin are shown for comparison. The correlation between the synaptic markers is high with no shift and it drops off sharply with lateral offset, as expected for tightly correlated presynaptic and postsynaptic markers. On the other hand, GAD and VGluT1 do not colocalize, because they label different synapse types, and the negative colocalization score with no shift gradually increases to 0 with lateral offset. D, Between layer differences in the astrocytic association of different synapse types.

In wild-type mice, we found that the majority of glutamatergic synapses (72±2%) are adjacent to astrocytic processes as detected by immunolabel to glutamine synthetase. This is very similar to previous EM estimates in mouse somatosensory cortex layer 4, where 68% of glutamatergic synapses on dendritic spines were in contact with astrocytic processes at the bouton-spine interface [69]. The proportion of glutamatergic synapses in contact with astrocytes was not uniform across layers, and we detected the highest association in layer 2/3 (Figure 7D), similarly to previous observations in rat visual cortex [70]. Interestingly, we observed significant differences in the astrocytic association of the different synapse types, as shown in Figure 7B. Thus, compared to the majority glutamatergic synapses containing VGluT1 (61±2% astrocytic association), significantly less VGluT2 synapses (46±1%, *p* < 0.001) were adjacent to astrocytes. The difference in astrocytic association was even more pronounced when considering the inhibitory GAD synapses which were half as likely to be adjacent to an astrocytic process (29 ± 1%, *p* < 0.001) compared to excitatory VGluT1 synapses.

Because there are no previous data about differences in glial association of cortical excitatory and inhibitory synapses, we sought additional evidence to confirm this finding. We quantified the colocalization between the astrocytic marker (GS) and the postsynaptic markers for excitatory (PSD95) and inhibitory (gephyrin) synapses using the van Steensel method [51]. This method evaluates the extent of spatial correlation by testing for the effect of very small relative displacements between pairs of marker images on a measurement of image overlap, the Pearson correlation coefficient. Because of the abundance of synaptic and glial markers, overlapping spatial distributions might occur by chance. If the association between two channels is real, however, then any shift of one channel relative to the other will decrease the observed degree of colocalization. On the other hand, if two channels tend to be mutually exclusive, then a shift will increase the degree of colocalization. Finally, if the association between two channels is occurring by chance, then a shift will not substantially affect the degree of colocalization. Consistent with our findings with the probabilistic synapse detector, the van Steensel method detected a colocalization between the astrocytic marker GS and the postsynaptic excitatory marker PSD95 7C. The degree of colocalization increased with shifts between the two channels of up to 0.3 −0.4*µm*, suggesting that the two markers are adjacent, but not overlapping. No colocalization was detected between GS and the postsynaptic inhibitory marker gephyrin. This does not mean that GS was not present next to a portion of inhibitory synapses, it only signifies that GS was not preferentially associated with inhibitory synapses. This result is consistent with our finding that excitatory synapses are more likely to be associated with astrocytic processes compared to inhibitory synapses.

Comparison of WT with KO mice revealed a number of significant changes in astrocytic involvement at synapses. Consistent with the detected overall changes in synaptic density, there were significant decreases in the densities of synapses adjacent to astrocytes, for almost all synapse types and sizes as shown in Figure 8A and in more detail in Supplemental Figure S5. The only exception were the small glutamatergic synapses in layer 4, for which the density of synapses adjacent to astrocytes increased in KO mice. While the density of synapses adjacent to astrocytes is very much influenced by the changes in overall synaptic density, the fraction of synapses adjacent to astrocytes reflects the actual changes in glial involvement in Fragile X Syndrome. A significant decrease in the fraction of synapses adjacent to astrocytes was detected for glutamatergic and VGluT1 synapses of all sizes, except small VGluT1 synapses (layers 1 - 4 combined) as shown in Figure 8B and in more detail in Supplemental Figure S6. VGluT2 synapses (medium size) and GAD (medium and large) also tended to decrease. When analyzed by cortical layer, there were significant decreases in the fraction of astrocytic association for large glutamatergic synapses in layers 1 and 2/3, large VGluT1 synapses in layer 1, small and medium VGLuT2 synapses in layers 2/3, and medium GAD synapses in layer 1. Interestingly, no changes in the astrocytic association for any of the synapse types and sizes were detected in layer 4.

**Figure 8.**
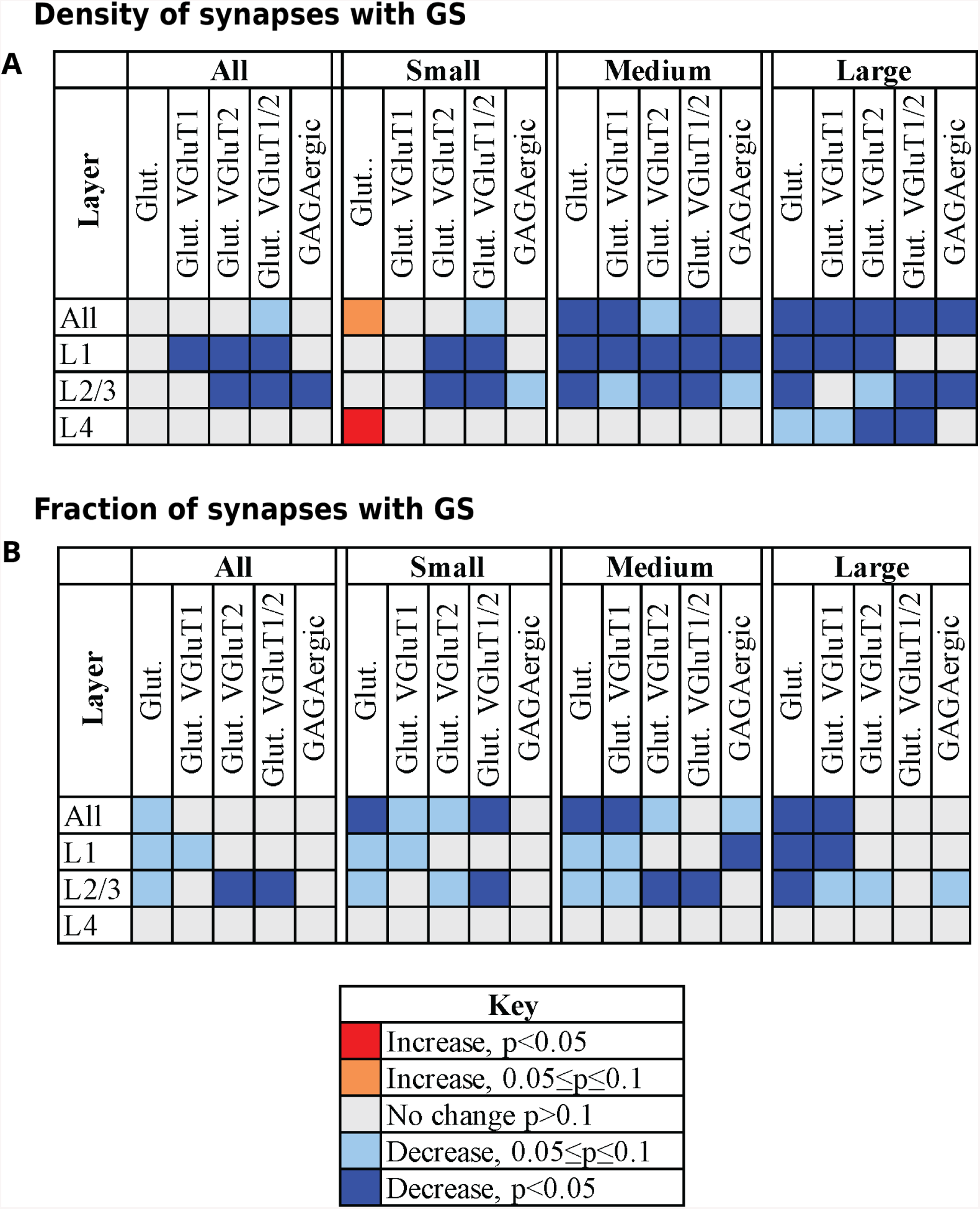
Summary of the astrocytic synapse density differences in the knockout mice. A, Changes in the density of synapses adjacent to astrocytic processes. The density decreases between the wild-type and knock-out mice for Layers 1-3 and the difference is especially pronounced for medium and large synapses. B, Changes in the fraction of synapses adjacent to astrocytic processes.

## Discussion

Using immunofluorescent array tomography and automatic probabilistic synapse detection methods we show wide-ranging changes of synapses and their association with astrocytes in the somatosensory cortex of adult FMR1 knock-out mice, a Fragile X mouse model. Overall, there is a significant decrease in the density of excitatory glutamatergic synapses and their association with astrocytes. However, the changes vary greatly, and are at times in opposite directions, depending on synapse type, size, as well as cortical layer. The changes in supragranular layers (layers 1 and 2/3) reflect the overall decrease in the density of glutamatergic synapses, both VGluT1 and VGluT2 type. Meanwhile in the granular layer (layer 4) there is a significant increase in the density of glutamatergic synapses, mostly due to an increase in small VGluT1 synapses, and no significant change in VGluT2 synapses. No changes in the astrocytic association of synapses are seen in layer 4, while in the supragranular layers a significantly lower fraction of glutamatergic VGluT1 synapses are associated with astrocytes. As for the inhibitory GABAergic synapses, the only change detected is an overall decrease in the density of large synapses, and a decrease in the astrocytic association of medium-sized synapses in layer 1. Overall, these changes result in an increased Excitation/Inhibition ratio in FMR1 knock-out mice. Thus, the absence of FMRP markedly alters the neocortical synaptic circuitry by both changing the relative contributions of synapses of different types, and the astrocytic involvement at synapses.

Our results are consistent with reported data in adult FMR1 knock-out mice suggesting cortical circuit changes such as increase in smaller size immature synapses and decrease in larger size synapses (as evidenced by the spine size in [35, 71]), as well as decreased association of astrocytes with hippocampal synapses [35]. We have extended these observations, by showing that these changes are not uniform, but depend on the synapse type, as well as cortical layer. A previous study had indeed shown that layers 4 and 5 synapses of different types in mouse somatosensory cortex exhibit various deficits in FMR1 knock-out mice [31] and we have now characterized the synapse type specific changes in the supragranular layers as well. Finally, we are showing for the first time that the changes in astrocytic involvement at synapses specifically affect excitatory glutamatergic synapses, and to a much lesser extent inhibitory GABAergic synapses in supragranular layers of mouse somatosensory cortex, with no detectable effect on layer 4 synapses.

A potential drawback of our study is the small number of analyzed animals, due to the laborious nature of these experiments. In order to minimize variations between experimental sessions and obtain more accurate results, our original design was to immunolabel and image the samples in pairs, one KO and one WT. The subsequent analysis revealed good consistency from experiment to experiment, as evidenced by the individual data points presented in the Supplemental Figures, which enabled the detection of multiple synaptic changes in the FMR1 KO mice regardless of the small number of animals. In addition, comparison of our results from WT mice to the available published data, further confirmed that we are not only able to correctly quantify the densities of the two main synapse types, excitatory and inhibitory, but that we are also detecting the known layer variations in VGluT2 synapses [63, 64, 65], and even the rather subtle layer variations in astrocytic association of excitatory synapses [70].

Previous studies of Fragile X syndrome have mostly focused on the alterations occurring in the synaptic circuitry, but we now show that there are specific deficits in cortical astrocytes and their interactions with synapses. Interestingly, these changes affect almost exclusively glutamatergic synapses. Indeed, it has been shown that glutamatergic but not GABAergic neurons, critically depend on the presence of glia to establish synaptic transmission [72]. Furthermore, studies employing selective deletion of FMRP in astrocytes strongly suggest the involvement of these glial cells in Fragile X pathogenesis (Hodges et al., 2017), likely through impaired glutamate uptake [73]. It thus appears that in Fragile X Syndrome astrocytes may mediate at least some of the pathological effects on glutamatergic synapses, while GABAergic synapses are likely influenced by a different mechanism.

Overall, our study reveals complex, synapse-type and layer specific changes in the somatosensory cortex of FMR1 knock-out mice. Some of these changes are in opposite directions, or affect only a small population of synapses and therefore become obscured when analyzing the overall synaptic content. The ability to dissect the deficits by specific synapse categories, as well as astrocytic involvement, are crucial for understanding the overall picture of synaptic changes, to begin to unravel the multiple ways in which they affect circuit function, and ultimately define targets for therapeutic treatment and prevention.

## Acknowledgments

This work was supported by the National Institutes of Health grants: NIH-TRA 1R01NS092474, R01 MH109475, R01 MH104227, and R01 NS094499. It was also supported by the following: National Science Foundation (NSF), United States Office of Naval Research (ONR), United States Army Research Office (ARO), and the National Geospatial-Intelligence Agency (NGA). Gifts to GS from Google, Microsoft, and Amazon are also acknowledged. The authors thank Prof. Richard Weinberg from UNC for his ever sharp insights about synapses.

## Supplemental tables

**Table S1.**
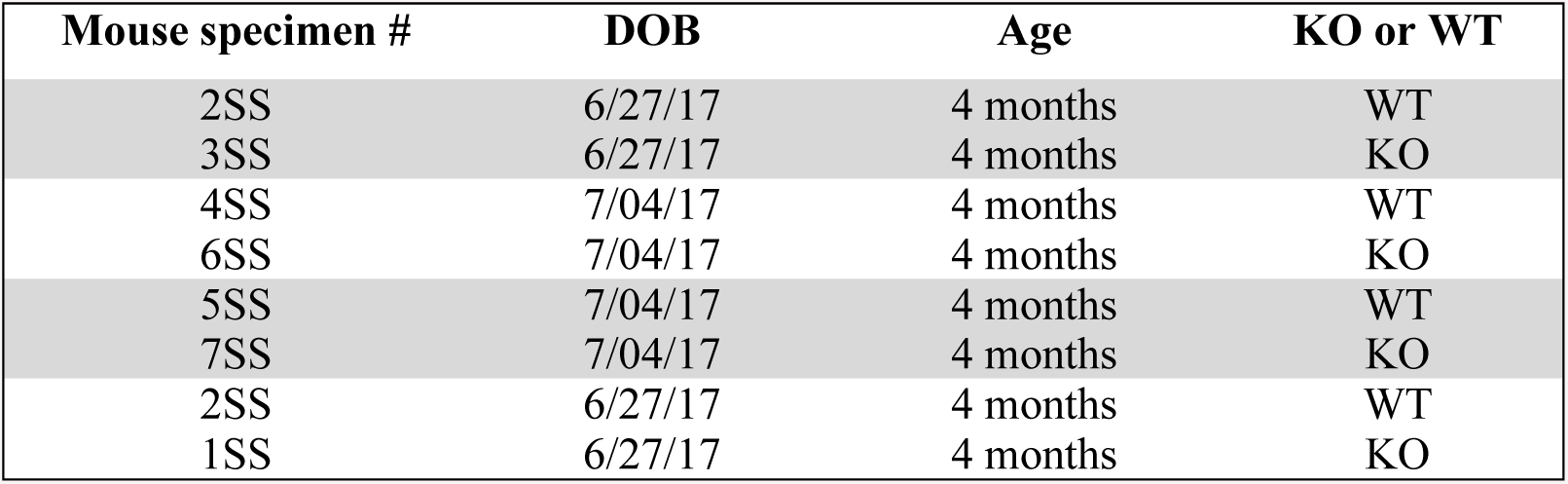
Mice used for the experiments and their condition. ‘WT’ refers to wild-type, ‘KO’ refers to knockout.

**Table S2.**
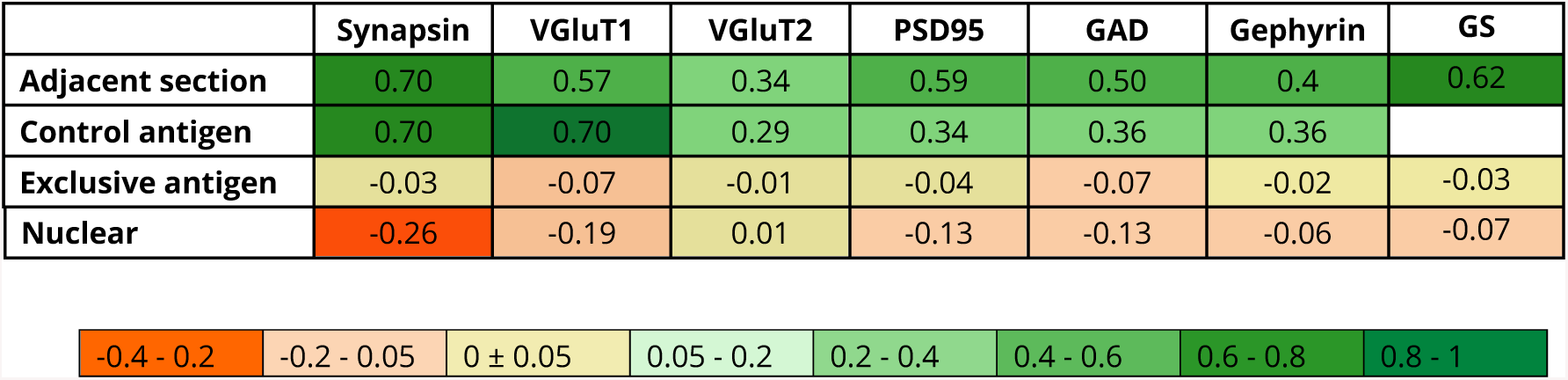
Antibody controls. Pearson’s correlation (PC) coefficients from 4 different control experiments are shown. *The comparison between adjacent sections* tests the consistency of staining, as the distribution of targets is very similar on two adjacent ultrathin sections (70 nm thickness). This correlation is influenced by antibody characteristics, but also the size of targets, with smaller targets displaying larger spatial variability from section to section. The comparison with an antibody against a *control* antigen is a test for the specificity of staining. The antibody staining pattern was compared to that for a control antigen with a similar distribution (overlapping or adjacent). The following comparisons were done: Synapsin/VGluT1 (overlapping), VGluT2/Synapsin (partially overlapping; only a subset of synapsin puncta are expected to overlap with VGluT2), PSD95/Synapsin (adjacent), GAD/Synapsin (partially overlapping; only a subset of synapsin puncta are expected to overlap with GAD), Gephyrin/GAD (adjacent). Control antigen was not available for GS. Another test for specificity is the comparison with an antibody against a spatially *exclusive antigen*. PC coefficient values of 0 and below are expected in this case. Synapsin (presynaptic protein) was compared with GS (astrocytic protein). VGluT1, VGluT2 and PSD95 (present in excitatory synapses) were each compared with GAD (in inhibitory synapses); GAD and gephyrin (inhibitory synapses) were compared with VGluT1 (excitatory synapses). And finally, all antibodies were compared with DAPI to control for background *nuclear* staining. All antibodies performed as expected, exhibiting good consistency of label between sections, strong colocalization with antibodies against control antigens, and negative colocalization with antibodies to exclusive antigens and the nuclear label DAPI. The only exception was the VGluT2 antibody, which had higher background label as confirmed by its scores. For this reason, the query for VGluT2 synapses was more stringent, requiring the presence of VGluT2 on 2 consecutive sections.

## Supplemental figures

**Figure S1.**
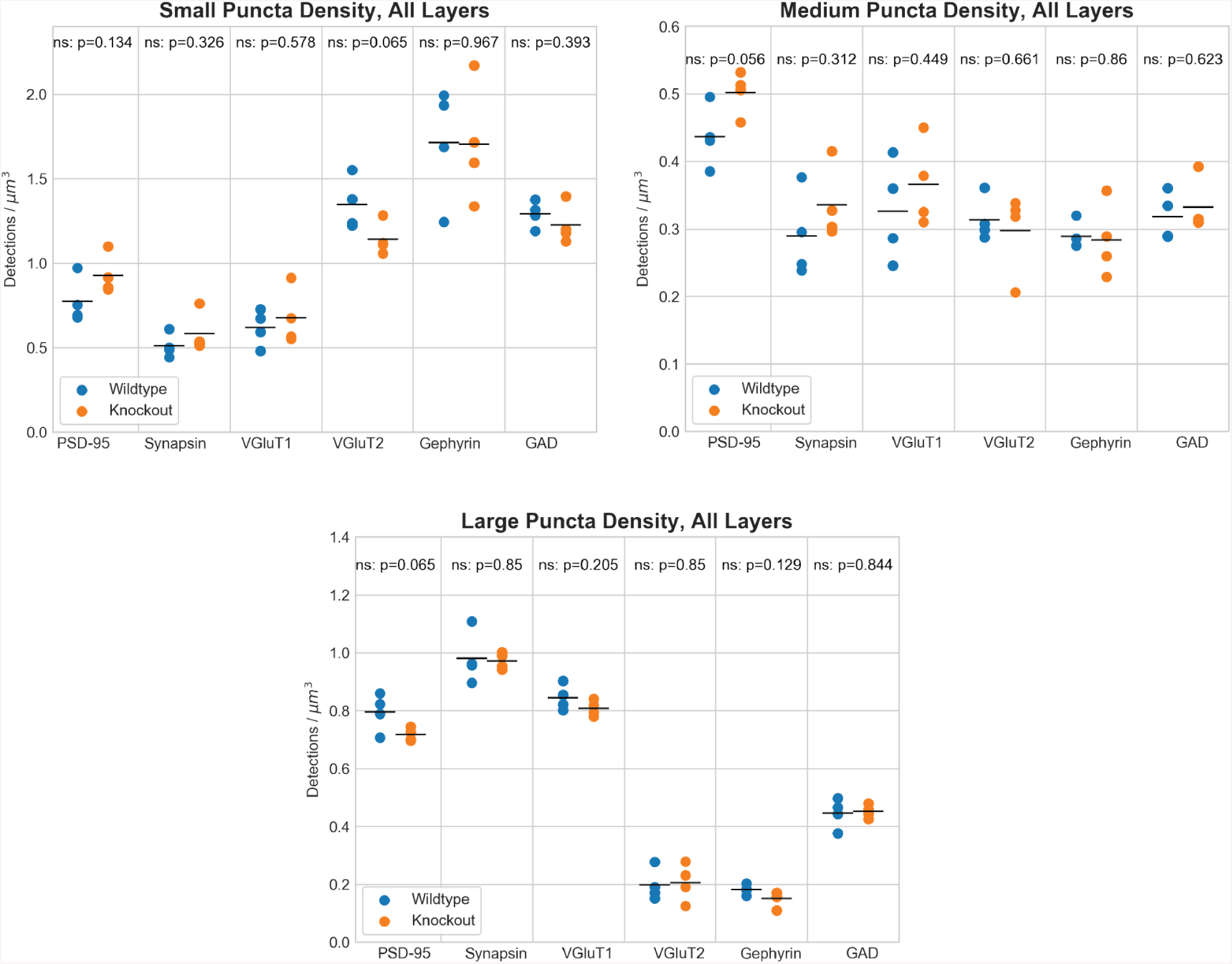
Density distribution of puncta from different synaptic markers, organized by size.

**Figure S2.**
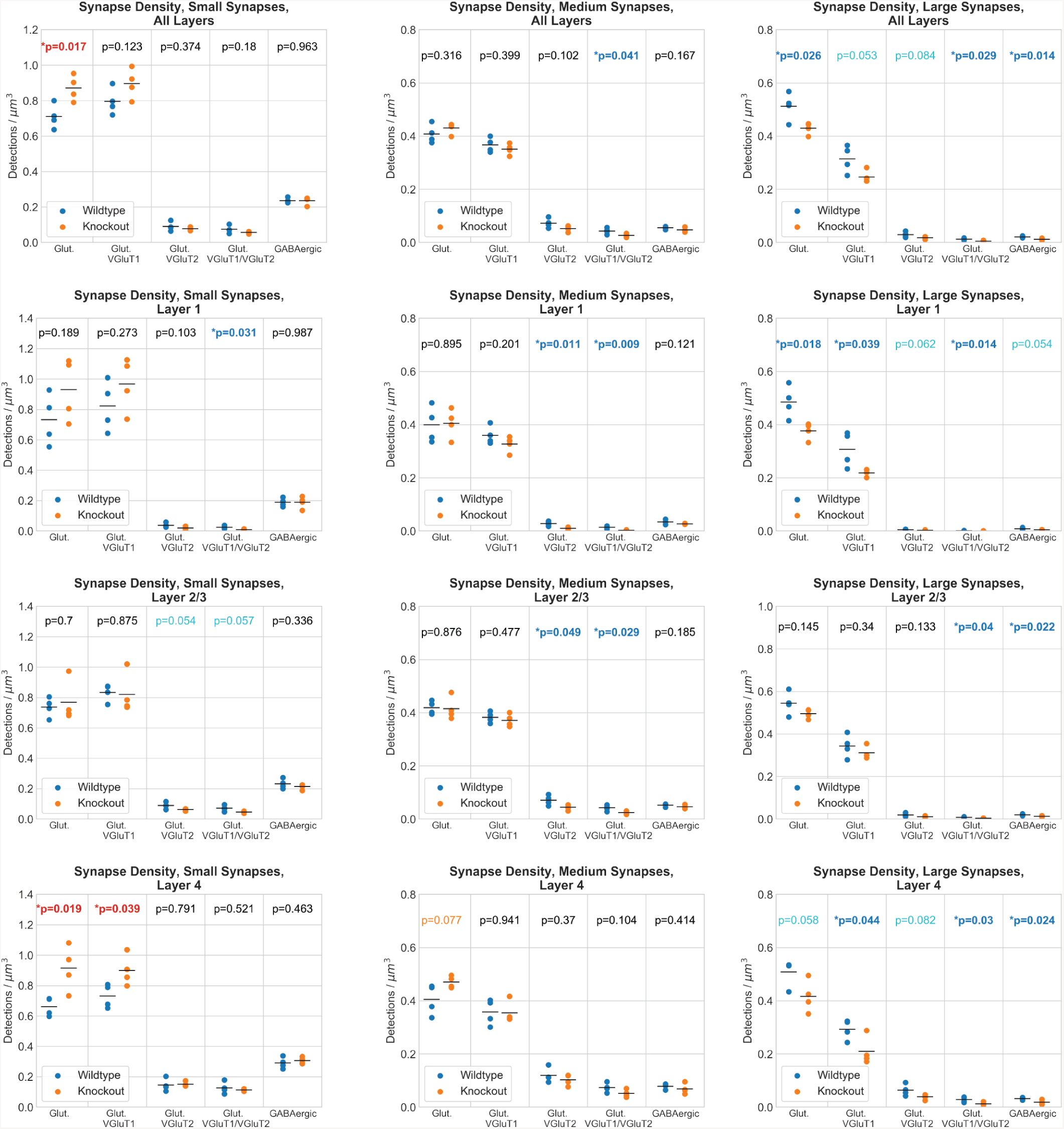
Synapse density distribution differences between wild-type and knockout mice. Organized by size, layer, and synapse type.

**Figure S3.**
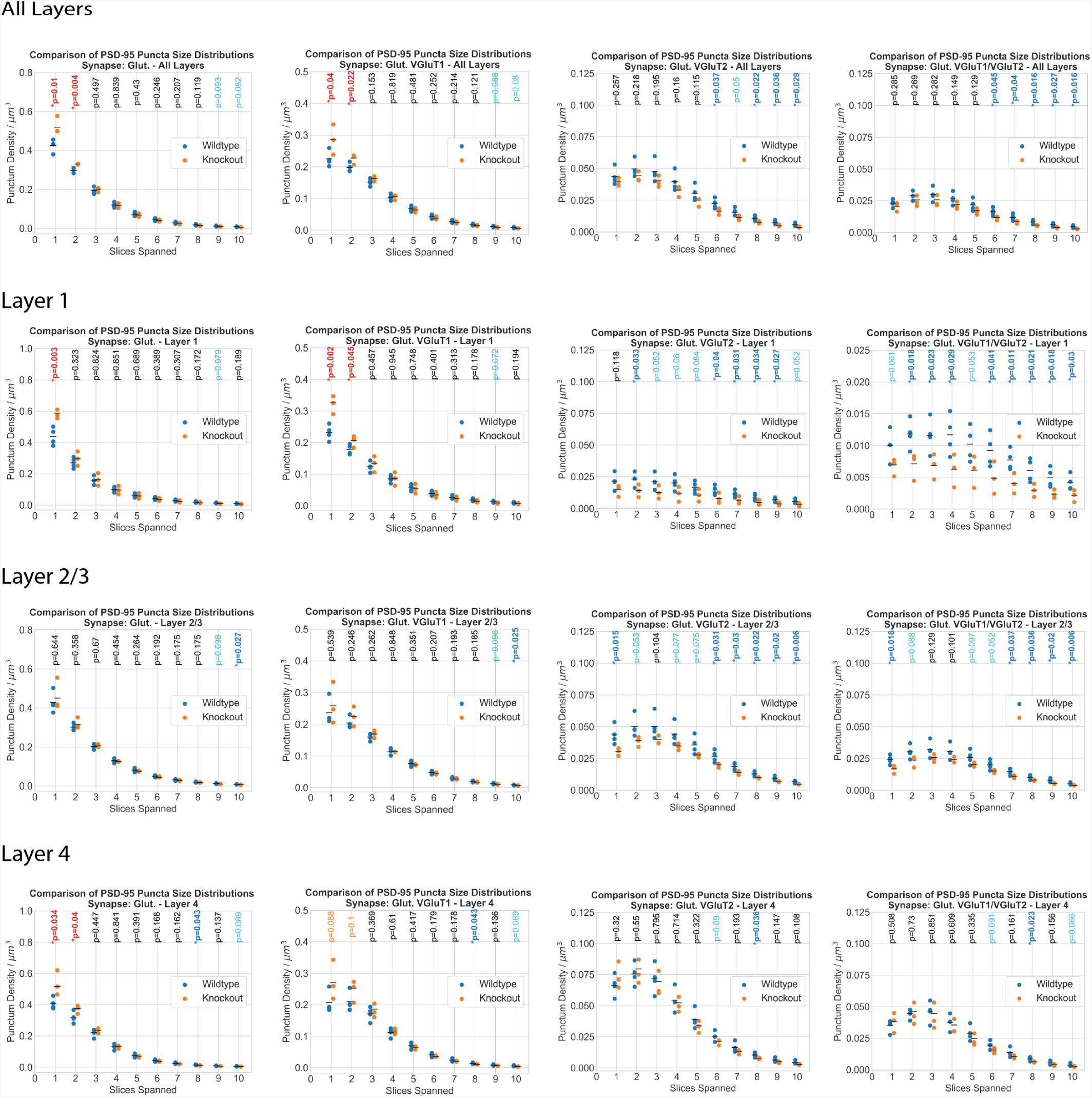
Histogram set showing the distribution of PSD-95 puncta associated with a synapse. Organized by size, layer, and synapse type.

**Figure S4.**
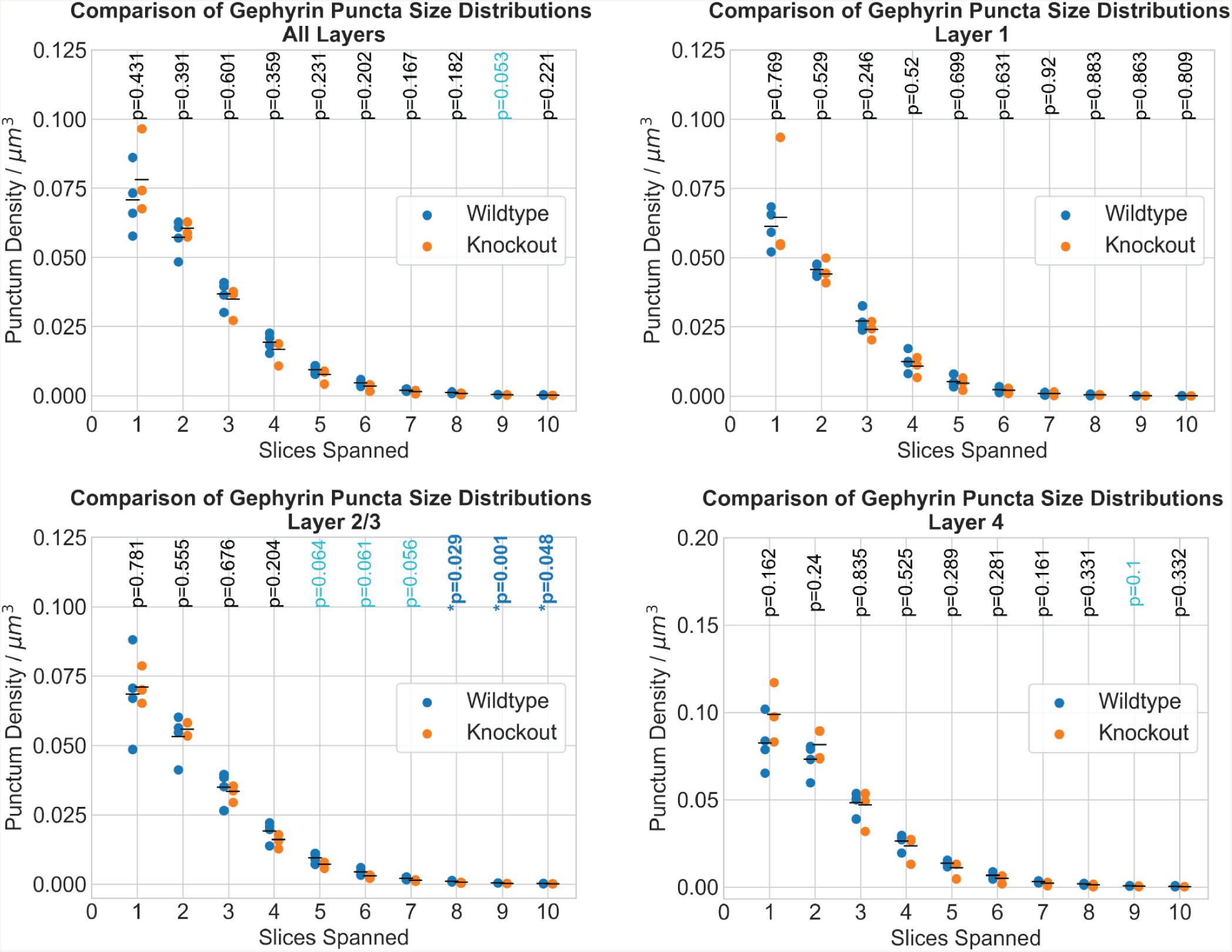
Histogram set showing the distribution of gephyrin puncta associated with a synapse. Organized by size, layer, and synapse type.

**Figure S5.**
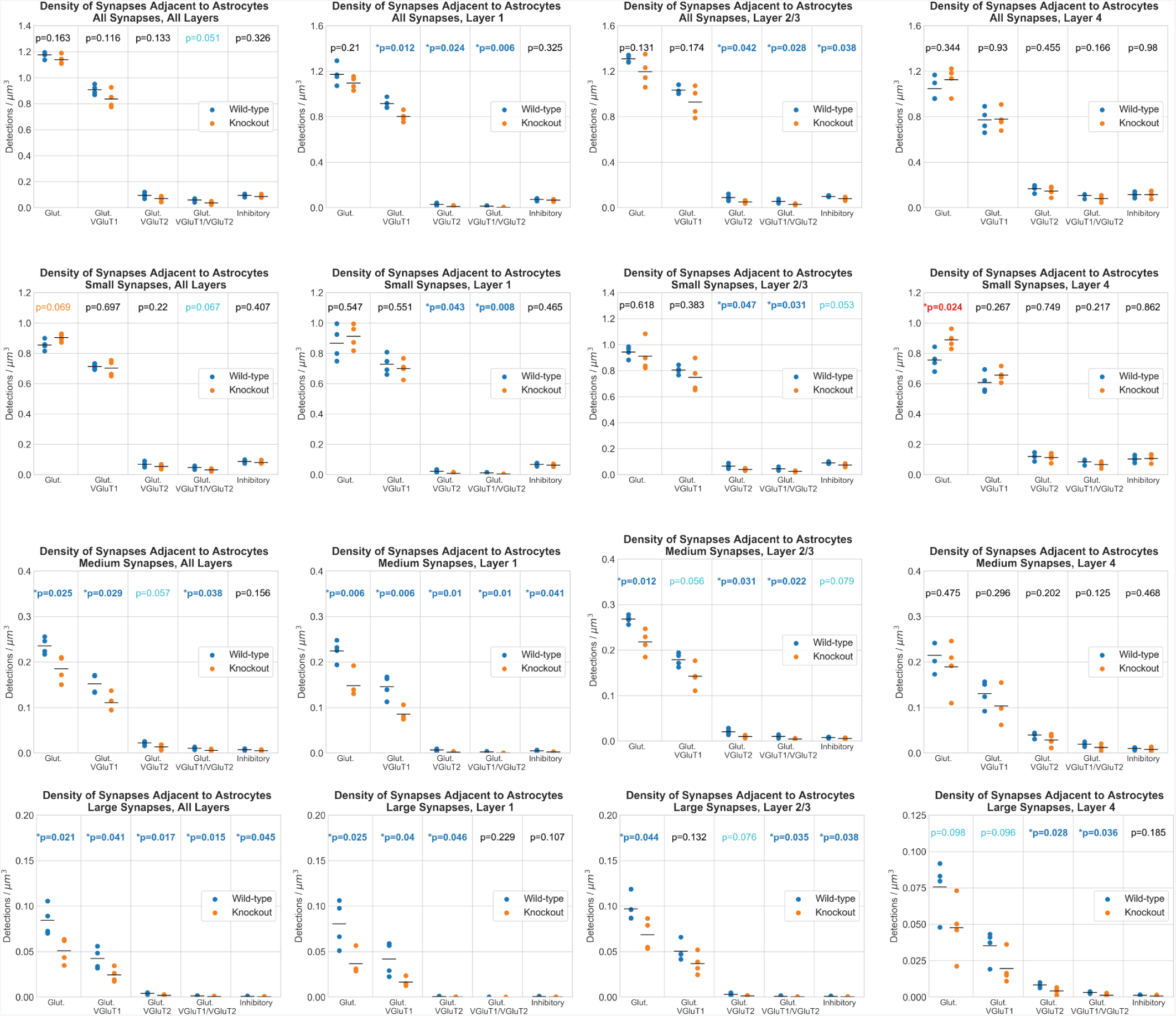
Comparison of the density of synapses associated with astrocytes between wild-type and knockout mice. Organized by size, layer, and synapse type.

**Figure S6.**
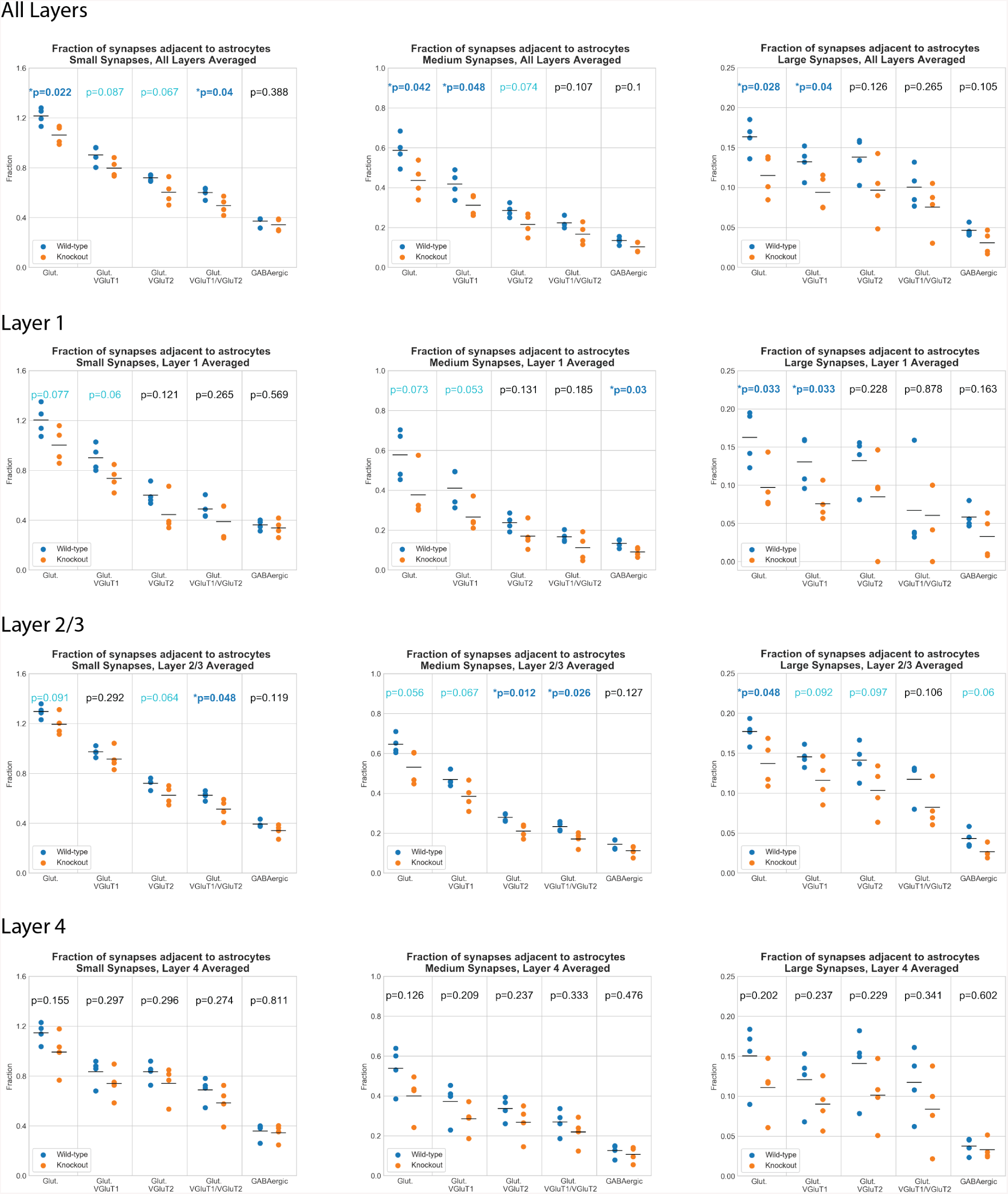
Fraction of astrocytic synapses to synapses between wild-type and knockout mice. Organized by size, layer, and synapse type.

